# Generation of a New Frizzled 2 Flox Mouse Model to Clarify Its Role in Development

**DOI:** 10.1101/2021.01.27.428341

**Authors:** Megan N. Michalski, Cassandra R. Diegel, Zhendong A. Zhong, Kelly Suino-Powell, Levi Blazer, Jarrett Adams, VAI Vivarium and Transgenics Core, Ian Beddows, Karsten Melcher, Sachdev S. Sidhu, Stephane Angers, Bart O. Williams

## Abstract

It is currently accepted that Wnt receptors, Frizzleds (Fzd), have high functional redundancy, making individual receptors challenging to target therapeutically. Specifically, Fzd2 is believed to be functionally redundant with Fzd1 and Fzd7, findings which were based largely on previously published global knockout mouse studies. Conversely, a Fzd2 global knockout mouse model developed by the International Mouse Phenotype Consortium (IMPC) is early embryonic lethal, suggesting Fzd2 is critical for early embryonic development. If global deletion of Fzd2 leads to early lethality, floxed models are necessary to identify tissue-specific phenotypes. We found that a previously published Fzd2 flox model does not fully delete Fzd2 function. To reconcile the contradictory findings in Fzd2 mouse models and allow for tissue-specific studies of Fzd2, we have generated a new flox model using a modified two-cell homologous recombination CRISPR approach. We demonstrated successful simultaneous insertion of two *loxP* sites fully surrounding the Fzd2 gene and confirmed cre-mediated recombination deletes the sequence between the *loxP* sites leading to a Fzd2 null allele. Preliminary studies suggest global knockouts are early embryonic lethal and full characterization of the tissue-specific effects of Fzd2 deletion is currently underway. This work suggests Fzd2 uniquely regulates development and emphasizes the importance of thorough validation of newly generated mouse models.

## Introduction

The use of genetically engineered mouse models has been crucial in the understanding of human disease. Conditional ablation of genes using the Cre-lox system was a significant advance in mouse modeling by allowing for tissue-specific deletion of target genes [1, 2]. This is particularly important in studying genetic modifications that cause lethality when deleted globally. While some reports indicate that simultaneous insertion of two *loxP* sites into the genome is possible with pronuclear injection of reagents to facilitate CRISPR/Cas9-mediated targeted insertions [3, 4], we have had challenges achieving this in our studies. We have found that floxed allele creation via CRISPR/Cas9-mediated methods typically requires a two-step approach. The first *loxP* site is incorporated using CRISPR/Cas9 methods and identifying founders with the desired insertion. This founder line is then expanded to allow for the second round of pronuclear microinjections to facilitate insertion of the second *loxP* site. This approach takes about one year to generate a mouse with a fully floxed allele [5]. Horii et al. sped up the process by introducing the two *loxP* sites with two sequential injections at the 1-cell and 2-cell embryonic stages before implantation [6]. A potential limitation of this method is that injecting the same embryo twice could lead to decreased zygote survival. Work done by Janet Rossant’s laboratory increased the efficiency of incorporating large insertions using CRISPR/Cas9 editing by injecting at the 2-cell stage [7]. They further increased efficiency by generating a monomeric streptavidin (mSA) tagged Cas9 mRNA and injected it with biotinylated PCR templates to localize the repair template to specific double-strand breaks. We modified this method to quickly and efficiently simultaneously insert two *loxP* sites surrounding the coding sequence for Frizzled 2 (Fzd2), a Wnt ligand-receptor. We chose to generate this floxed allele to overcome unanticipated genomic alterations we discovered in previously generated mouse models of Fzd2.

Our interests in Wnt signaling regulation of development and disease led us to specifically study Fzd2 due to the association of *FZD2* heterozygous mutations with the human syndromes Autosomal Dominant Omodysplasia (ADO) [8-11] and Robinow Syndrome (RS) [12]. These syndromes present with limb reductions and craniofacial anomalies. It is unclear how Fzd2 regulates these structures’ development and how the human mutations alter Fzd2 function. Mouse models targeting Fzd2 can help address these questions. Through extensive investigation of previously published Fzd2 mouse models, we have determined the available models do not fully delete Fzd2 protein or function. We find evidence that a previously published Fzd2 knockout animal (*Fzd2*^*tm1*.*1Nat*^) [13, 14] is an allele with hypomorphic function. While mice homozygous for this allele do not express protein containing the N-terminus of FZD2, we noted that the *Fzd2*^*tm1*.*1Nat*^ allele preserves a portion of the sequence encoding the FZD2 C-terminus, which retains signaling function. Mice homozygous for the *Fzd2*^*tm1*.*1Nat*^ allele progress normally through embryonic development and develop partially penetrant cleft palate (CP) (∼50 %).

In contrast to the *Fzd2*^*tm1*.*1Nat*^ allele, the International Mouse Phenotyping Consortium (IMPC) generated a global null allele of *Fzd2*, which deletes the entire coding region and homozygosity leads to early embryonic lethality (before E9.5) [15]. Because of this early embryonic lethal phenotype, floxed alleles of *Fzd2* are necessary to evaluate FZD2 function in adult and later-stage embryonic tissues. We obtained a previously reported *Fzd2* flox strain [16], but we determined that complete deletion of Fzd2 did not occur after Cre exposure.

Given our interest in Wnt signaling at the plasma membrane and the unanticipated complications we found with the currently available Fzd2 mouse models, we generated a new Fzd2 flox mouse model to help delineate the roles of Fzd2 in development. Here, we report a method to generate a floxed mouse line in one step using a modified two-cell homologous recombination (2C-HR)-CRISPR protocol [7]. Additionally, we aim to encourage the regular practice of whole-genome sequencing (WGS) mouse models to verify off-target modifications or other unintended alterations.

## Materials and Methods

### Experimental animals

The mice used in this study were maintained in accordance with institutional animal care and use guidelines. Experimental protocols were approved by the Institutional Animal Care and Use Committee of the Van Andel Institute. Mice were fed LabDiet 5021 mouse breeder diet and housed in Thoren Maxi-Miser IVC caging systems with a 12-hour light/12-hour dark cycle. The *Fzd2*^*tm1*.*1Nat*^ mice [13, 14] were obtained from the laboratory of Jeremy Nathans (Johns Hopkins University) and the *Fzd2*^*tm1Eem*^ flox mice [16] from the laboratory of Edward Morrisey (University of Pennsylvania). Our newly generated B6;C3-*Fzd2*^*tm1Vari*^ flox allele will be referred to as *Fzd2*^*tm1Vari*^.

### Guide RNA design for inserting *loxP* sites

Briefly, guide RNAs (gRNA) targeting the promoter region of Fzd2 and downstream of the 3’ UTR were designed using the CRISPOR website (http://crispor.tefor.net/) against *Mus musculus* (GCA_001632555.1) genome reference sequence [17] and were purchased from Integrated DNA Technologies (IDT). We used sgRNA AGGGAGCAGCTTCGCCAGTT to target the promoter region and sgRNA GAGCCGGGTTCATTAAAGTT to target downstream of the 3’UTR.

### pNS20-SpCas9-mSA Construct and Recombinant Protein Generation and Purification

To generate pNS20-SpCas9-mSA, we PCR amplifying the optimized linker and mSA coding sequence from the pCS+Cas9-mSA plasmid, created by the laboratory of Janet Rossant (Addgene, 103882) [7]. A BamHI site was incorporated to the 5’ end of the mSA fragment and a MluI site to the 3’ end. We modified the original BamHI site present at the 3’ end of the mSA fragment from a GGA to a GGT codon to eliminate the restriction site but retain the glycine. BamHI and MluI restriction sites were used to remove the SNAP coding sequence and insert the mSA coding sequence into pNS20-SpCas9-SNAP (Laboratory of Gerald Schwank, Addgene, 113717) [18]. Recombinant protein was generated and purified as previously described [19]. Briefly, N-terminally His6-MBP-tagged Cas9 mSA was expressed in *E. coli* Rosetta (DE3) pLysS cells (Novagen) in 6L of LB medium to an OD_600_ of ∼1 at 30 °C and induced with 100 μM isopropyl-β-D-thio-galactopyranoside (IPTG) at 16 °C overnight. For purification of the Cas9 fusion protein, cells were harvested, resuspended, and lysed in 20 mM Tris, pH 8.0, 500 mM NaCl, 5 mM imidazole, using a French Press with pressure set to 900 Pa. Lysates were cleared by centrifugation for 1 hour at 38,700 x g and passed over a 5 mL HisTrap HP column (GE Healthcare). The column was washed with 15 column volumes of 20 mM Tris, pH 8.0, 500 mM NaCl, and 5 mM imidazole and then eluted with 20 mM Tris, pH 8.0, 250 mM NaCl, 250 mM imidazole. To the eluted protein, 1 mg of TEV protease was added for each 25 mg of Cas9 and dialyzed together against 20 mM HEPES, pH 7.5, 100 mM KCl at 4 °C overnight. The next morning the cleaved protein was loaded on a 5 ml HiTrap HeparinHP column (GE Healthcare). The protein was eluted over a gradient from 100 mM to 2M KCl. The protein was further purified by size-exclusion chromatography through a HiLoad 26/60 Superdex 200 column (GE Healthcare) in 20 mM HEPES, pH 7.5, 250 mM KCl. DTT was added to 2 mM and glycerol to 10% to the final purified protein for storage.

### In vitro DNA cleavage by recombinant Cas9-mSA protein

The DNA cleavage activity of our Cas9-mSA endonuclease was assayed on linearized plasmid DNA containing eGFP. The plasmid was digested overnight with XbaI followed by column purification (28104, Qiagen). We used RNA guide GTGAACCGCATCGAGCTGAA to target eGFP [20] which was complexed in vitro with our Cas9-mSA enzyme. The cleavage assay was performed as previously described [21]. In short, the cleavage reaction contained the following components: 1 ul sgRNA (1 ug/ul), 3ul 10X Cas9 nuclease reaction buffer (1M NaCl, 0.1M MgCl_2_, 0.5M Tris-HCl, 1mg/mL BSA, pH= 7.9), 600 ng of either wild-type Alt-R™S.p Cas9 nuclease (1081058, IDT) or S.p Cas9-mSA protein, 500 ng of linearized plasmid, and ddH_2_O (to a final volume of 30 µl). The mixtures were incubated at 37 °C for 2 hours, after which 1 μl Proteinase K (20 mg/ml) was added, and the mixture was subsequently incubated at 65 °C for 10 minutes to release the DNA from the Cas9 protein. Mixtures lacking the sgRNA targeting eGFP were used as a negative control.

### BIO-PCR donor template

Double-stranded DNA fragments with ∼1kb spanning homology arms around a *loxP* sequence were synthesized as gBlocks Gene Fragments (IDT). The donor template had a disrupted PAM sequence and a unique XhoI restriction site in the 5’ donor template and an AgeI restriction site in the 3’ donor template. 5’ biotin-modified oligos were used to PCR amplify the biotinylated template, which then underwent PCR cleanup and ethanol precipitation before microinjection [7].

### Mouse lines and embryos

Embryos were generated by intercross of B6C3F1/J animals (JAX stock #100010). Thirty 3-4-week-old females were superovulated by intraperitoneal (IP) injection of pregnant mare serum gonadotropin (5 IU) followed by IP injection of human chorionic gonadotropin (5IU) 48 hours later. Mating was established on the same day. The B6C3F2 two-cell embryos used for injection were collected at two different time points, with half recovered at 0.5 days post coitum (dpc) and the other half at 1.5 dpc. Those recovered at 0.5 dpc were cultured for 24 hours in K-RVCL (Cook Medical), while embryos collected at 1.5 dpc were placed in K-RVCL and microinjected within 6 hours after recovery.

### Microinjection and embryo transfer

Microinjection commenced immediately following recovery of the 1.5 dpc embryos. A modified Integrated DNA Technologies (IDT, https://www.idtdna.com/pages) mouse zygote microinjection of Alt-R™ CRISPR-Cas9 System ribonucleoprotein delivery protocol was used. Complete sgRNA’s from IDT (crRNA and tracrRNA) were incubated with Cas9-mSA protein to generate ribonucleoprotein (RNP) complexes before the biotinylated template (BIO-template) was added [22]. Both complexes were combined for a final microinjection mix containing 12.5 ng/ul sgRNA: Cas9-mSA protein, 4 ng/ul BIO-template targeting both the 5’ and 3’ regions in injection buffer (1 mM Tris-HCl, pH 7.5; 0.1 mM EDTA). RNP complexes were injected into each of the two-cells nuclei under negative capacitance generated via MICRO-ePORE pinpoint cell penetrator (World Precision Instruments). Embryos were microinjected in paraffin oil-covered M2 (Cytospring) and then transferred to K-RVCL for culture. Twenty-four hours later, 12-15 of these ∼8-cell embryos were transferred unilaterally into pseudopregnant females. Founders were identified by standard PCR methods and backcrossed to wild-type C57BL/6J once before intercrossing to generate animals for our study.

### Genotyping by PCR and indel sequencing

Genomic DNA from mouse tail biopsies was isolated by alkaline digestion. To genotype the *Fzd2*^*tm1Vari*^ flox 5’ *loxP* allele we used the following primers: Fzd2-5’*loxP*-Fwd (GAGATTACAGGTGTGAGCTACTG) and Fzd2-5’*loxP*-Rev (CTGGAGAGGGAAGGGAATTTG) to amplify a 367bp wild type product and a 398bp *loxP* product. To genotype the 3’*loxP* allele we used the following primers: Fzd2-3’*loxP*-Fwd (GAGAAAGCAGGAGGATGTAAGAG) and Fzd2-3’*loxP*-Rev (CTGTCCCACCTTCCATCAAAT) to amplify a 292bp wildtype product and a 324bp *loxP* product. To determine Cre mediated recombination both in vitro and in vivo, we used the following primers: Fzd2-5’*loxP*-Fwd, Fzd2-3’*loxP*-Fwd, and Fzd2-3’*loxP*-Rev to amplify a 485bp knockout product, a 324bp *loxP* product, and a 292bp wild type product. All PCR products for each model were Sanger sequenced by GENEWIZ (https://www.genewiz.com/en) for validation and to confirm correct *loxP* insertion. SnapGene software (from Insightful Science; available at snapgene.com) was used for sequence alignment and animal model design.

To genotype the *Fzd2*^*tm1Eem*^ flox allele, we used the following primers as previously described [16]: Fzd2-Flox_F (GCCTGCTCGCTATTTTTGTTGGC) and Fzd2-Flox_R (AAATGAGGAGGGAGAAAGAGGGGG) to amplify a 210bp wild type and 300bp *loxP* product.

### Generation of KO alleles

To generate global knockout animals, male mice homozygous for the *Fzd2*^*tm1Vari*^ floxed allele were bred to female B6.C-Tg(CMV-cre)1Cgn/J (JAX stock #006054; [2]). The resulting offspring were genotyped to determine allele-specific Cre-mediated recombination for Fzd2 using the following primers: Fzd2-5’*loxP*-Fwd (GAGATTACAGGTGTGAGCTACTG) and Fzd2-3’*loxP*-Rev (CTGTCCCACCTTCCATCAAAT) to amplify a 485bp knockout product. PCR products were Sanger sequenced by GENEWIZ (https://www.genewiz.com/en) to validate correct PCR products, indel insertion, and recombination.

Similarly, we generated global mutants for the *Fzd2*^*tm1Eem*^ allele. To determine Cre mediated recombination within this allele, we designed a primer upstream of the 5’ *loxP* site within the 5’ UTR. We used the following primers for determining Cre recombination: Fzd2-AS-F (AAACCGACTAATTGGGATCGG) along with Fzd2-Flox_F (GCCTGCTCGCTATTTTTGTTGGC) and Fzd2-Flox_R (AAATGAGGAGGGAGAAAGAGGGGG) R to amplify a 474bp knockout, 210bp wild type, and 300bp *loxP* product.

### Restriction Digestion of PCR Products

PCR products for the 5’ *loxP*, 3’ *loxP*, and Cre mediated KO recombination allele were amplified from the *Fzd2*^*tm1Vari*^ flox model, column purified (T1030S, New England Biolabs) and then used for a restriction endonuclease digest. Digestion reactions consisted of 500ng of PCR product, 20 units of either XhoI (R1046S, NEB) or AgeI (R3552S, NEB) in their specified buffers. The reactions were incubated at 37 °C for 1 hour before samples were run on a 2% agarose gel for imaging.

### Whole Genome Sequencing (WGS) -Illumina and ONT long-read sequencing

Genomic DNA (gDNA) was extracted from whole blood via cardiac puncture from 6-8-week-old *Fzd2*^*tm1Eem*^ (homozygous flox or homozygous global mutants created by crossing with CMV-cre mice) or *Fzd2*^*tm1Vari*^ (homozygous flox) animals using Quick-DNA™ Miniprep Plus Kit (Zymo Research). Illumina libraries were constructed with average insert sizes of ∼800 bp and sequenced on an Illumina NovaSeq6000 using paired-end 150bp reads, and an average of 30x coverage was achieved. The resulting reads were aligned (with BWA-MEM aligner) to mm10 mouse reference genome for sequence and structural accuracy. Nanopore long-read sequencing was performed using the same *Fzd2*^*tm1Eem*^ homozygous flox/flox genomic DNA used for Illumina sequencing on a MinION flow cell (Oxford Nanopore Technologies). The resulting reads were aligned (with Minimap2 aligner) to mm10 mouse reference genome. The resulting BAM files were visualized with Integrative Genomics Viewer (The Broad Institute).

### Limb micromass cultures

Limbs were harvested from E11.5-E12.5 *Fzd2*^*tm1Eem*^ or *Fzd2*^*tm1Vari*^ embryos for micromass culture according to previously published protocols [23, 24] with modifications. Briefly, fore- and hindlimbs were removed from embryos and digested in ∼10U/mL of Dispase II (Gibco) in DMEM (Gibco) at 37 °C for 1 hour. Yolk sacs were collected for genotyping. Limbs were further dissociated by pipetting up and down and then centrifuged at 300 x g for 5 minutes. Cell pellets were resuspended in culture medium (DMEM, 4.5 g/L glucose, 1 mM sodium pyruvate, 2 mM L-glutamine, 10 % FBS, 1 % PenStrep) and passed through a 40um cell strainer. Cells were centrifuged at 300 x g for 5 minutes and resuspended in culture medium to reach a cell concentration of 10-20×10^6^ cells/mL. Cells were plated in a 10ul micromass drop in a 24-well tissue culture well and incubated at 5 % CO_2_, 37 °C tissue culture incubator for 2 hours to allow cell attachment followed by the addition of 1mL of culture medium. The following day, the cells were gently washed with 1X PBS and incubated for 1 hour with Ad5CMV:eGFP or Ad5CMV:Cre-eGFP (MOI 100, University of Iowa Viral Vector Core Facility). Culture media was added to inactivate viral complexes. Cells were imaged for eGFP incorporation 48 hours later, and gDNA was collected for Sanger Sequencing (GENEWIZ; https://www.genewiz.com/en) or cell suspensions for flow cytometric analyses. Alternatively, RNA was extracted from micromass cultures for qPCR 72 hours after Ad-Cre transduction.

### Flow cytometry

Single-cell suspensions were generated from limb micromass cultures or from limb buds dissected from E12.5 embryos for flow cytometric analyses. Limb micromass cultures were trypsinized in TrypLE (Gibco) and neutralized with media containing FBS. Following centrifugation at 300 x g for 5 minutes, cells were resuspended in cold flow buffer (1X HBSS without magnesium, calcium, or phenol red, with 2 % FBS), and cells were filtered through a pre-wet 40um cell strainer into a 50mL conical tube. Single-cell suspensions from limb buds were prepared following a modified protocol [25, 26]. Briefly, limb buds from E12.5 mouse embryos were dissected and digested in 1 ml 1X HBSS (without magnesium, calcium, or phenol red) containing 1 mg/ml collagenase D (Millipore Sigma) and 50 µg/ml DNase I (Roche) at 37 °C for approximately 45 minutes. Limb buds were gently pipetted every 5 minutes with a 1000 uL pipette tip until the tissue was dissociated into a single-cell suspension. Cold flow buffer was added to stop the digestion, and the cell suspension was filtered through a pre-wet 40um cell strainer into a 50mL conical tube and centrifuged at 300 x g for 10 minutes. Approximately 2 × 10^5^ (micromass) or 1 × 10^6^ (limb bud digestion) cells were stained with 50 nM unconjugated Fzd2 antibody (methods for antibody generation are detailed in [27]) for 30 minutes on ice. Cells were washed and stained with anti-F(ab’)2-647 (1:1000, Jackson ImmunoResearch) for 30 minutes on ice. Cells were washed and resuspended in 200uL flow buffer containing 5ug/mL DAPI prior to analyses. Alternatively, if cells were fixed prior to analysis, they were stained with Zombie Violet™ Fixable Viability Kit (1:1000, BioLegend) for 30 minutes on ice during secondary antibody staining. Cells were then washed and fixed in 4 % paraformaldehyde (PFA) ice for 20 minutes, washed, and resuspended in 200uL of flow buffer. Cells were analyzed on CytoFLEX S (Beckman Coulter).

### Quantitative real-time polymerase chain reaction (qRT-PCR)

RNA was extracted from *Fzd2*^*tm1Eem*^ micromass cultures using Quick-RNA Miniprep kit (Zymo Research) followed by cDNA conversion using High-Capacity cDNA Reverse Transcription Kit (Applied Biosystems). Real-time quantitative PCR was performed with SYBR green master mix (Life Technologies) on StepOnePlus™ Real-Time PCR System (Applied Biosystems). Two sets of Fzd2 primers were used targeting either the Fzd2 CDS region or 3’ untranslated region (UTR) that is upstream of the *loxP* insertion site in both Fzd2-flox models. mFzd2-CDS-F GCTGCGCTTCCACTTTCTTC; mFzd2-CDS-R GAACGAAGCCCGCAATGTAG; mFzd2-3UTR-F GCACCCTACGGACTCCTATT; mFzd2-3UTR-R GGTGTTGGGCAGAGTTCTTT.

For copy number detection, gDNA was harvested from limb buds from E14.5 *Fzd2*^*tm1*.*1Nat*^ or lung tissue from *Fzd2*^*tm1Eem*^ 8-week-old mice using Quick-DNA™ Miniprep Plus Kit (Zymo Research). A modified PCR protocol was used [28]. Briefly, the genomic DNA was diluted to ~30 ng/uL, and 2 uL was used in a 20 uL PCR reaction. Fzd2 Cq values were normalized to Fzd1, which was considered as an internal reference. The primers used were the same as those for limb bud qPCR described above.

### *Fzd2*^*tm1*.*1Nat*^ C-terminal peptide studies

The truncated mouse Fzd2 gene (Fzd2-delN) was cloned from pRK5-mFzd2 (#42254, Addgene) with a 1D4 tag at the C terminus by PCR using the following primers. SalI-mFzd2-5del-F AATAGTCGACGCCATGGGCCAGATCGAC; NotI-1D4-R aagcagcggccgcTTAGGCAGGCGCCACTTG. The resulting plasmids were transfected into HEK293 or HEK293T-FZDless [29] (gift from Dr. Benoit Vanhollebeke, Université libre de Bruxelles, Belgium) expressing a Wnt reporter gene (super top flash, STF) with X-tremeGENE HP DNA Transfection Reagent (Roche). The luciferase activity was detected 48 hours after transfection using the luciferase assay system (Promega) and a BioTek Synergy Neo Microplate Reader (BioTek).

## Results

### Previous mouse models of Fzd2 do not produce null alleles

We obtained two previously published Fzd2 mouse models: the *Fzd2*^*tm1*.*1Nat*^ allele [13] which was reported to be a global knockout, and the *Fzd2*^*tm1Eem*^ allele [16] which was intended to be a conditional allele. However, we found neither of these two models fully deleted Fzd2. Mice homozygous for the *Fzd2*^*tm1*.*1Nat*^ allele do not express N-terminus FZD2 protein (**Figure 1A and 1C**), but we noted that the *Fzd2*^*tm1*.*1Nat*^ allele retains a portion of the CDS (531bp) that could retain partial function (**Figure 1A and 1B**). We detected mRNA expression of this C-terminal fragment in E11.5 limb lysates isolated from *Fzd2*^*tm1*.*1Nat*^ homozygotes (**Figure 1D**). There are several potential alternative start codons within the remaining sequence, including an in-frame ATG with a strong Kozak sequence [30] that is present near the 5’ end of this Fzd2 fragment (a 531bp in-frame CDS is predicted, **Figure 1A**). Because there are no antibodies available that recognize the C-terminal region of FZD2, we expressed a C-terminal, epitope-tagged (1D4) version of this *Fzd2* fragment (Fzd2-delta-N), which initiates transcription from this ATG in HEK293 and HEK293T-FZDless cells (**Figure 1E**). This fragment localizes similar to full-length FZD2 (**Figure 1F**) and can activate the β-catenin-responsive TOPFlash reporter in HEK293 cells in response to WNT3A and RSPO1 [31] (**Figure 1G**). Fzd2-delta-N activity depends on endogenous FZDs as it has no Wnt signaling activity in HEK293T-FZDless cells that lack all 10 FZDs (**Figure 1G**) [29]. One possibility is that the carboxyl portion of FZD2 can partner with at least one wild-type Fzd family member to transduce canonical Wnt signaling. Another possibility is that FZD-delta-N can interact with other endogenous FZD-regulating proteins to affect other FZDs. Collectively, our studies support the hypothesis that the *Fzd2*^*tm1*.*1Nat*^ allele is hypomorphic.

**Figure 1.**
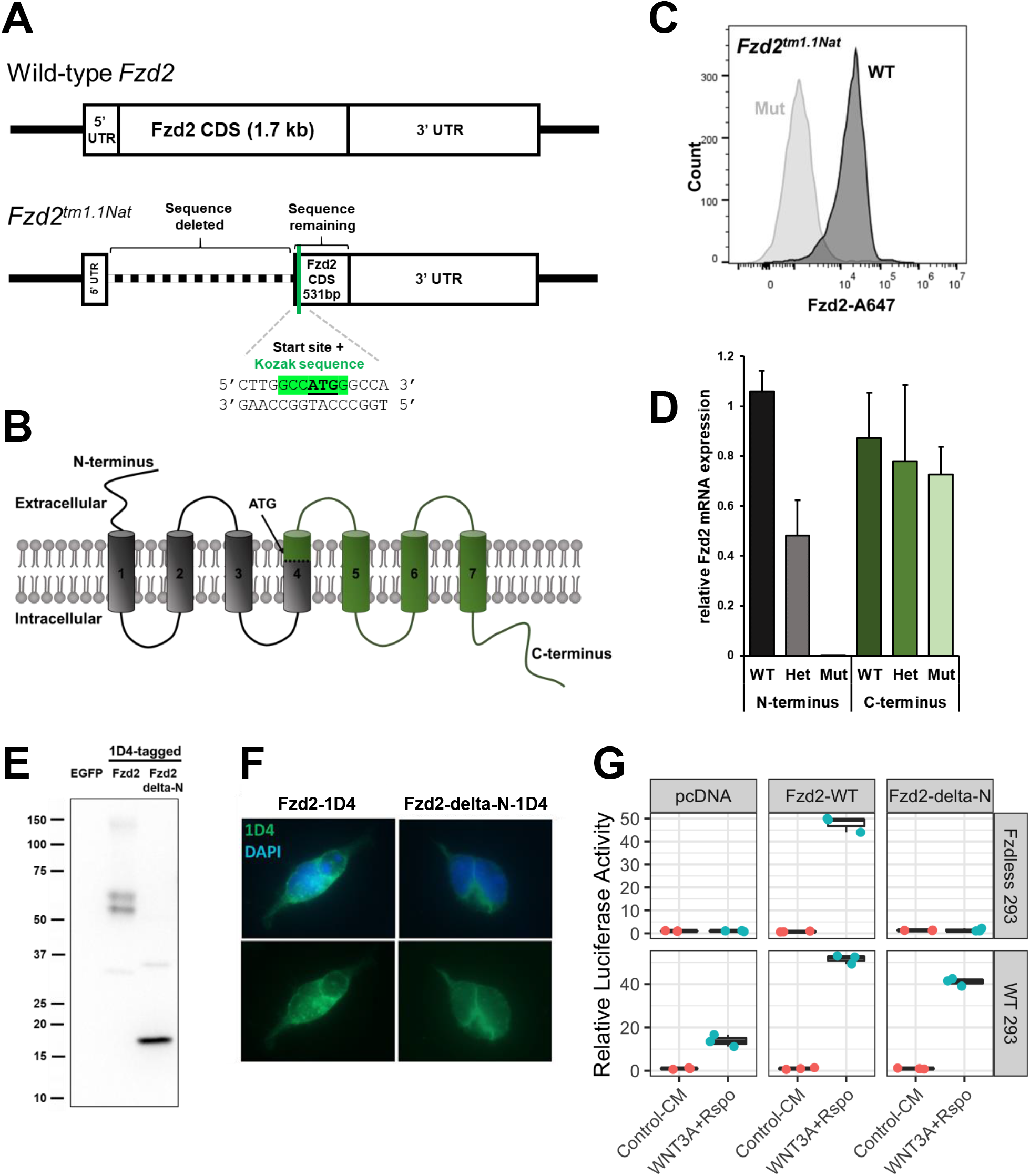
*Fzd2*^*tm1*.*1Nat*^ allele retains functional C-terminal peptide. A. *Fzd2*^*tm1*.*1Nat*^ allele retains 531bp of Fzd2 C-terminal sequence that could encode a protein with an in-frame ATG and Kozak sequence (green). **B**. Putative partial product with start site in the 4th transmembrane domain (green). **C**. Flow cytometry of E14.5 limb bud cells using antibody targeted to N-terminal FZD2. Cells were gated to discard debris and doublets, and live cells were determined by DAPI exclusion. FZD2 median fluorescent intensity (MFI) was compared between cells wild-type (WT) or homozygous (Mut) for the *Fzd2*^*tm1*.*1Nat*^ allele. WT MFI = 18294; Mut MFI = 1267 **D**. qRT-PCR of E14.5 limb buds from littermates wild-type (WT), heterozygous (Het), or homozygous (Mut) for the *Fzd2*^*tm1*.*1Nat*^ allele. The relative transcription of either the N-terminal (grey bars) or C-terminal (green bars) portions are compared. Note the C-terminal PCR primers are located in the remaining Fzd2 sequence (green transmembrane domain in panel B). **E**. A plasmid directing expression of the C-terminal fragment (Fzd2-delta-N) produces a stable protein when ectopically expressed in HEK293T cells (detected with a 1D4 tag). **E**. Immunofluorescence for the 1D4 epitope reveals similar localization of full-length Fzd2 (Fzd2-1D4) and the C terminal fragment (Fzd2-delta-N-1D4). **F**. Wild-type (WT) HEK293 cells or HEK293T cells lacking all 10 FZDs (FZD-less) were transfected with pcDNA-based plasmids expressing empty vector (pCDNA), Fzd2-WT or Fzd2-delta-N. Cells were treated with control conditioned media from L cells (Control-CM) or Wnt3A conditioned media from L cells and R-spondin1 conditioned media from HEK293T cells (Wnt3A+Rspo). Fzd2-WT increases luciferase activity from a β-catenin responsive reporter (TOPflash). Fzd2-delta-N increases luciferase in WT, but not FZD-less cells.

We obtained a previously reported *Fzd2*^*tm1Eem*^ flox strain [16] but determined that complete deletion of Fzd2 did not occur after exposure to Cre. WGS of mice homozygous for *Fzd2*^*tm1Eem*^ flox confirmed the allele is a complex genomic alteration consisting of a *Fzd2* gene duplication (**Figure 2A “flox” and 2B; Supplemental Figure 1A and 1B**). Complete details are indicated in the legend for Figure 2 and Discussion. Briefly, the *Fzd2*^*tm1Eem*^ flox allele contains two copies of Fzd2, which are separated by a large insertion. This insertion has 10kb of exogenous DNA in addition to a region (**Figure 2A**, black curved line) which is too long to be sequenced with a single continuous read. Additionally, we noted a 100 kb endogenous sequence downstream of the Fzd2 gene, which is duplicated and likely inverted (**Figure 2A and 2B**), but extensive analysis needs to be completed.

**Figure 2.**
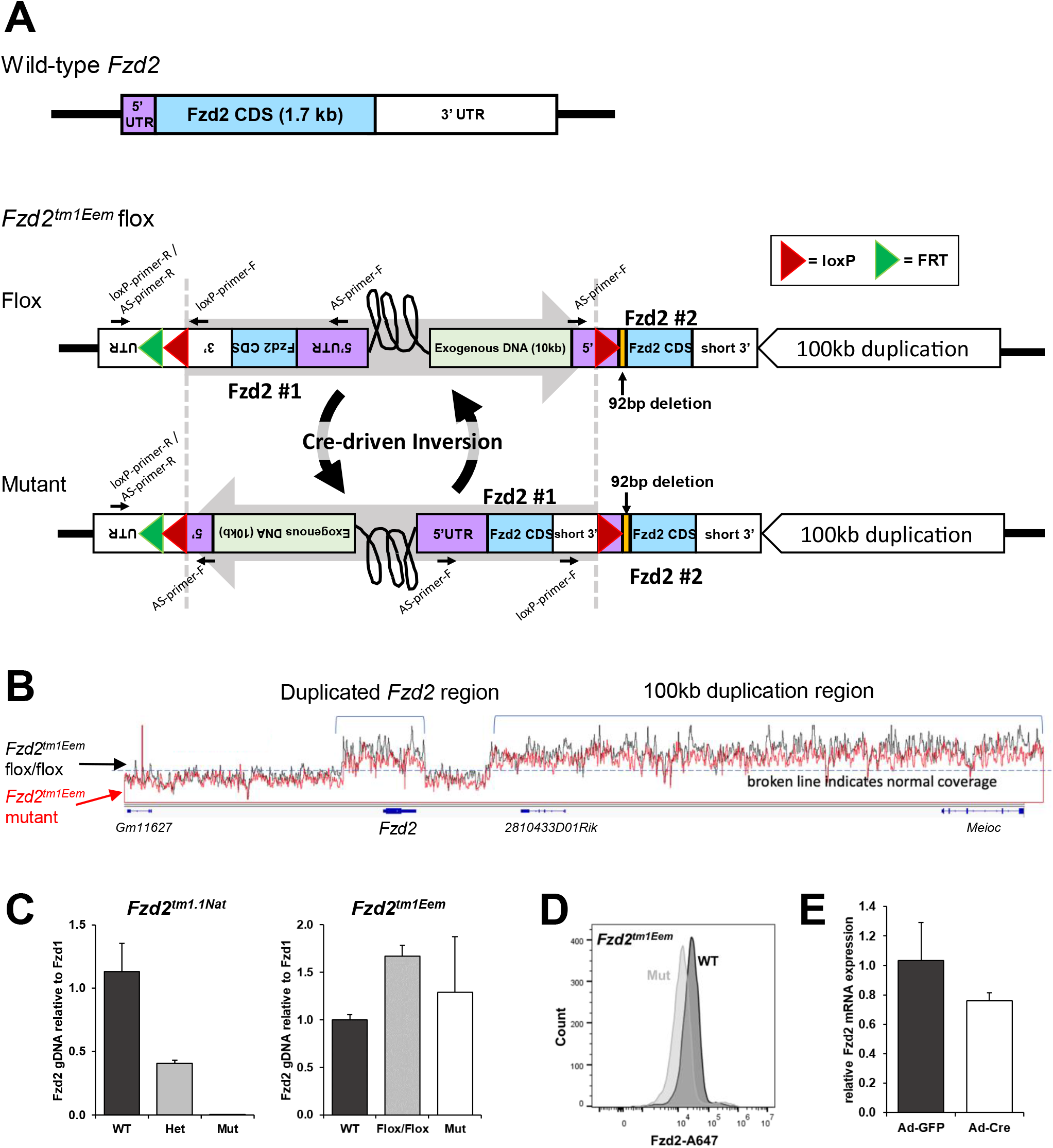
The published *Fzd2*^*tm1Eem*^ flox allele is a complex genomic alteration. A. Maps detailing our working model of the *Fzd2*^*tm1Eem*^ flox allele. A wild-type Fzd2 gene structure is shown, with 5’UTR in purple, CDS in blue, and 3’UTR in white. The *Fzd2*^*tm1Eem*^ flox allele contains a duplication of Fzd2, resulting in two copies (blue). There is 10 kb of exogenous DNA (bacterial gDNA and targeting plasmid PL253 sequence, light green) in between the two copies of Fzd2. “Fzd2 #1” contains the 3’ *loxP* insertion but not 5’*loxP*. “Fzd2 #2” contains 5’ *loxP* insertion as well as a 92bp deletion (orange) downstream of the 5’ *loxP* insertion site and immediately upstream of the Fzd2 start sequence. “Fzd2 #2” has a shortened 3’ UTR that is connected to an inverted 100 kb sequence downstream of the Fzd2 gene (refer to Supplemental Figure 1A for more details). The sequence between the two Fzd2 copies is of unknown length and denoted by a black curved lined (refer to Supplemental Figure 1B for more details). Because we cannot confirm the orientations of the two *loxP* sites within the two Fzd2 copies due to the long sequence between the two Fzd2 copies, we hypothesize cre-driven recombination results in continuous sequence inversion between the two *loxP* sites due to the opposite *loxP* orientations (refer to Discussion for more details). **B**. The IGV coverage tracks of *Fzd2*^*tm1Eem*^ flox/flox and homozygous global mutant WGS reads were merged to show no Fzd2 or other genomic DNA deletion after Cre exposure. **C**. Real-time quantitative PCR on genomic DNA shows a 50% decrease in Fzd2 gene content in *Fzd2*^*tm1*.*1Nat*^ heterozygotes and no Fzd2 gene content in homozygous mutants (primer targeting the N-terminal region). The *Fzd2*^*tm1Eem*^ display a ∼2-fold increase in Fzd2 gene content in flox/flox and ∼ homozygous mutant (Mut) samples, which supports Fzd2 being duplicated and *loxP* sites being in opposite orientations. **D**. FZD2 flow cytometric analysis of E14.5 limb buds from wild-type (WT) or mice homozygous for a germline cre-mediated recombination of the *Fzd2*^*tm1Eem*^ allele (Mut). Cells were gated to discard debris and doublets, and live cells were determined by DAPI exclusion. FZD2 median fluorescent intensity (MFI) was compared between WT and Mut. WT MFI = 25954; Mut MFI = 12680 **E**. Limb micromass cultures from homozygous flox *Fzd2*^*tm1Eem*^ mice were infected with Ad5CMV:eGFP (control) or Ad5CMV:Cre-eGFP at an MOI of 100 and assessed by qRT-PCR for Fzd2 mRNA expression.

One of the Fzd2 CDS regions is within the sequence flanked by *loxP* sites, which we believe are in opposite orientations. We hypothesize this leads to the sequence inverting between the *loxP* sites upon cre-mediated recombination, resulting in the two Fzd2 copies being in tandem (**Figure 2A “mutant”**). Consistent with the duplication seen via WGS, genomic Fzd2 DNA copy number was increased in homozygous flox mice (**Figure 2C, right**), and germline cre-mediated recombination only mildly reduced Fzd2 protein and mRNA expression in limb buds (**Figure 2D and 2E**). Due to the complex issues noted with the *Fzd2*^*tm1Eem*^ flox allele, we generated the *Fzd2*^*tm1Vari*^ flox allele using a modified approach [7].

### Successful simultaneous insertion of *loxP* sites using two-cell homologous recombination (2C-HR)-CRISPR

We cloned, expressed, and purified Cas9-streptavidin (Cas9-mSA) fusion protein (**Supplemental Figure 3A**) and confirmed that it cut linear plasmid similar to conventional Cas9 protein (**Supplemental Figure 3B**). To simultaneously incorporate *loxP* sites surrounding the Fzd2 gene, a mix contained two ribonucleoprotein complexes, each containing Cas9-mSA protein, sgRNA, and a biotinylated repair template targeting either the 5’ *loxP* or 3’ *loxP* were injected into 2-cell embryos (either 1-cell cultured to 2-cell, or 2-cell immediately after harvest) (**Supplemental Figure 3C**). To avoid altering endogenous Fzd2 expression, the 5’ *loxP* site was inserted into a region upstream of the promoter (**Figure 3A**). The 3’ *loxP* site is 994bp downstream of the 3’ UTR (**Figure 3A**). Of the 550 embryos microinjected, 314 were implanted. This produced 85 live-born animals, of which five genotyped for the 5’ modification and nine genotyped for the 3’ modification. Four out of 85 animals had both 5’ and 3’ modifications (**Supplemental Figure 4A and 4B**). The founders were all generated from the group collected as 1-cell zygotes, cultured, and then injected at the 2-cell stage. The PCR products were Sanger sequenced to confirm the *loxP* sequence and identify indels (**Figure 3B**). These founders were backcrossed to C57BL/6J animals to verify if the 5’ and 3’ *loxP* sites were inserted in cis (same allele) or trans (opposite alleles) and to produce offspring with germline mutations. Three of the four founders produced offspring with both the 5’ and 3’ *loxP* modifications (**Supplemental Figure 4C**). gDNA was harvested from wild-type or *Fzd2*^*tm1Vari*^ homozygous flox animals, and PCR amplified for either the 5’ *loxP* (**Figure 3C**) or the 3’ *loxP* (**Figure 3D**) product. The PCR products were purified and digested with XhoI (5’ *loxP*) or AgeI (3’ *loxP*), resulting in expected product sizes (**Figure 3C and 3D**).

**Figure 3.**
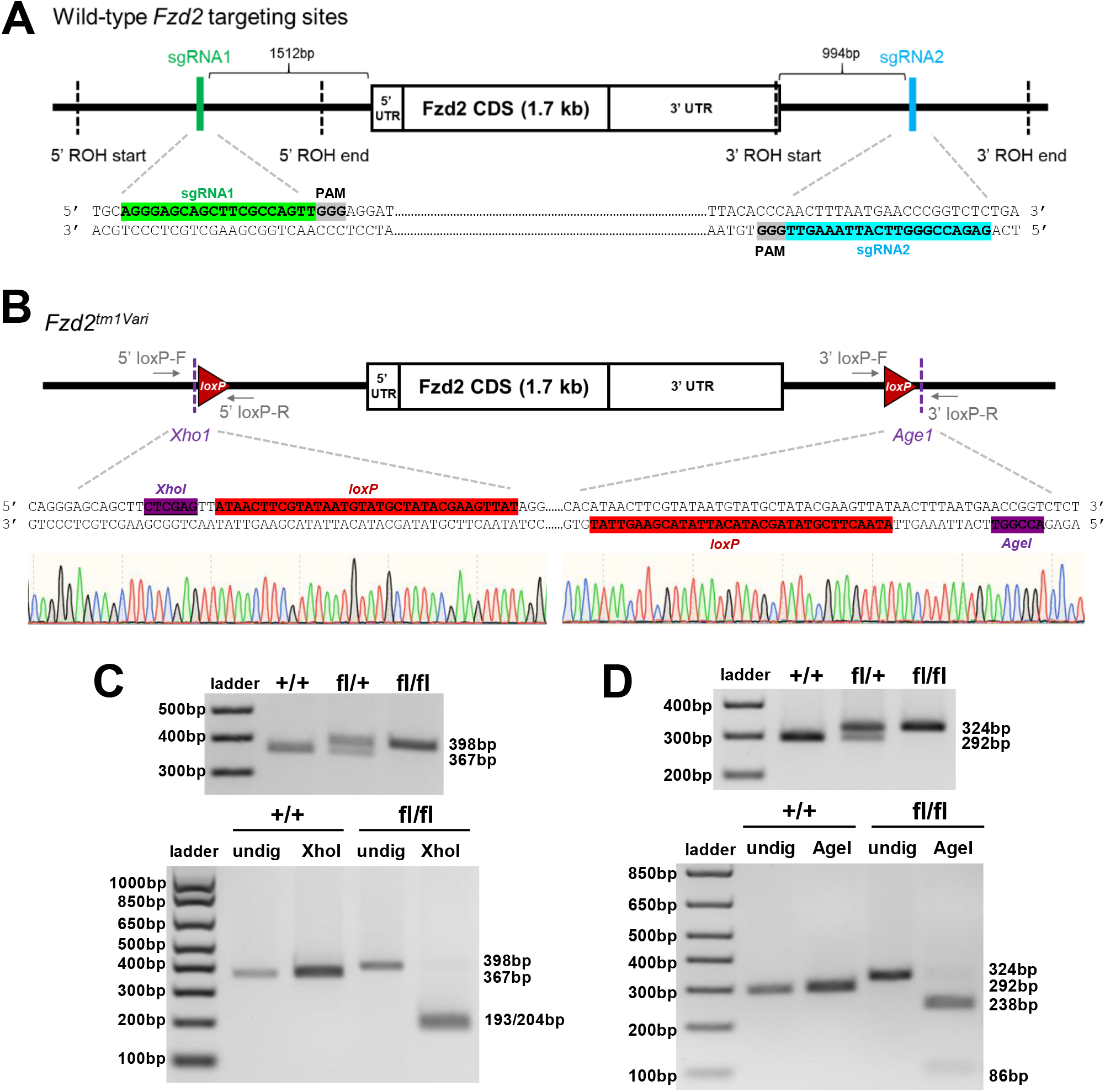
Generation and confirmation of *Fzd2*^*tm1Vari*^ allele. A. Map detailing the location of guide RNAs (indicated by the green and blue text on map and sequence) and regions of homology (ROH). **B**. Map of B6;C3H-*Fzd2*^*tm1Vari*^ intended modifications. *LoxP* sites were placed around the entire single exon *Fzd2* gene. Xho1 and Age1 restriction sites (purple) were incorporated to aid in validation of the allele. *LoxP* sites are indicated by red triangles, and 5’ and 3’ untranslated regions (UTR) are shown. Primer binding sites for genotyping are shown in grey. **C**. PCR-based genotyping products for 5’ *loxP* site (top) and digestion with XhoI of flox product (bottom) results in expected products (193bp and 204 bp). **D**. PCR-based genotyping products for 3’ *loxP* site and digestion with AgeI of flox product results in the expected products (86bp and 238 bp). undig = undigested

**Figure 4.**
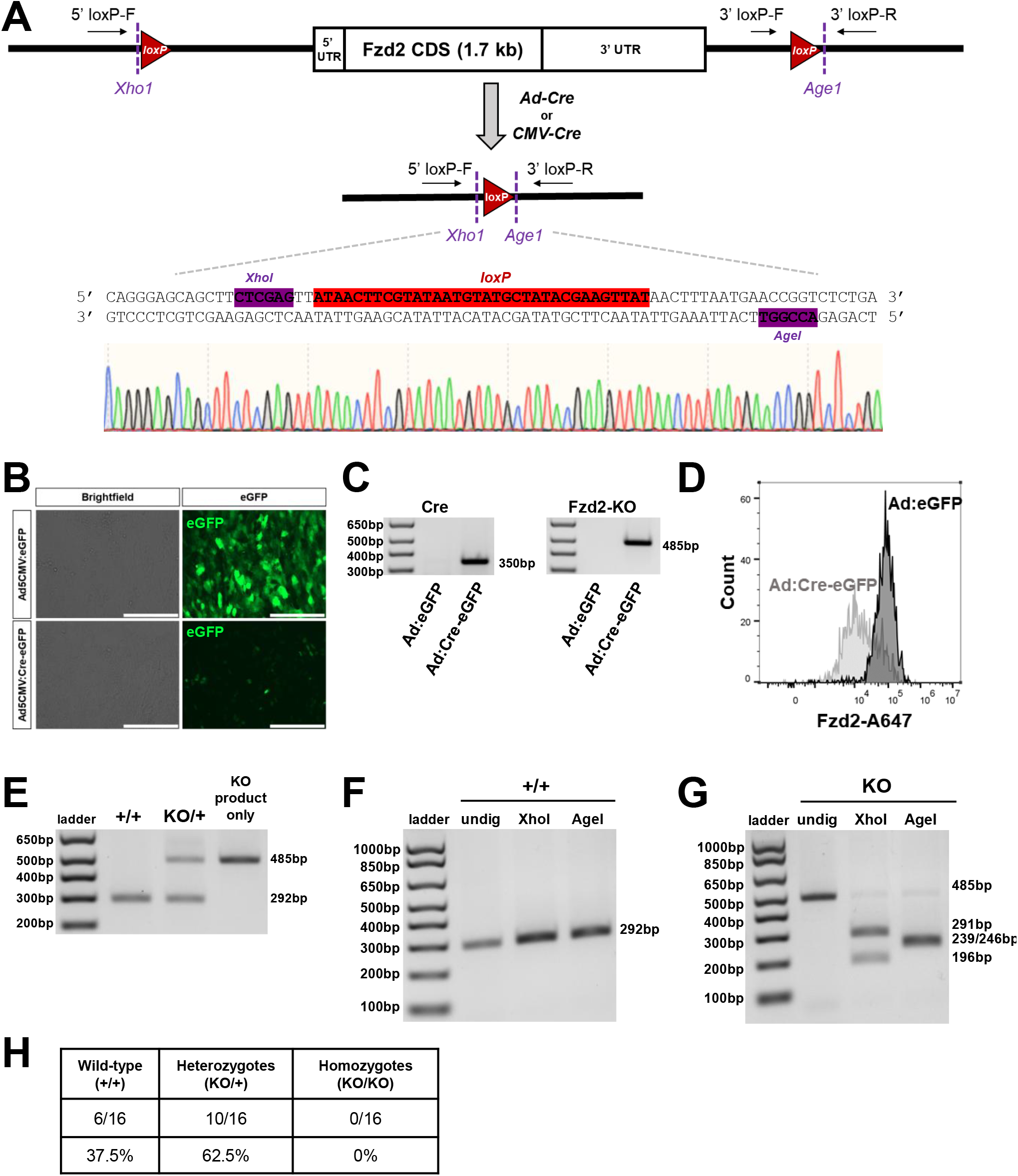
Confirmation of Cre-mediated recombination of *Fzd2*^*tm1Vari*^ allele. A. Map of proposed *Fzd2*^*tm1Vari*^ recombination product. Xho1 and Age1 restriction sites (purple) were incorporated to aid in the validation of the allele. *LoxP* sites are indicated by red triangles, and 5’and 3’ untranslated regions (UTR) are shown. Primer binding sites for genotyping are shown. **B**. Limb micromass cultures from *Fzd2*^*tmVari*^ flox/flox were infected with Ad5CMV:eGFP (control) or Ad5CMV:Cre-eGFP at an MOI of 100. Brightfield and fluorescent images show infection efficiency. Ad5CMV:eGFP is cytoplasmic and Ad5CMV:Cre-eGFP is nuclear. Scale = 200um **C**. PCR-based genotyping for Cre or Fzd2 knockout (KO). Ad5CMV:Cre-eGFP infected cells amplified a product for Cre and Fzd2-KO. **D**. Single-cell suspensions were prepared from Ad:eGFP or Ad:Cre-eGFP infected micromass cultures, stained for Fzd2, and analyzed by flow cytometry. Cells were gated to discard debris and doublets. Live cells were determined by DAPI exclusion followed by gating on eGFP. FZD2 median fluorescent intensity (MFI) was compared between Ad:eGFP or Ad:Cre-eGFP infected micromass cultures. **E**. PCR-based allele-specific genotyping products for wild-type (292bp) and KO (485bp) products. The third lane includes a PCR product using primer set for KO only to be used for restriction digestions. **F**. Wild-type product was digested with XhoI and AgeI resulting in a wild-type uncut product (292bp). **G**. KO product was digested with XhoI and AgeI resulting in the expected digestion products: XhoI (196 and 291bp), AgeI (239 and 246bp). undig = undigested

To further confirm the modifications produced in the newly generated *Fzd2*^*tm1Vari*^ founders were correct, gDNA was collected from a mouse homozygous for the floxed allele from one of the founders and whole-genome sequenced using traditional short-read Illumina sequencing (**Supplemental Figure 2**). The aligned reads confirmed the 5’ and 3’ *loxP* sites were in the correct location and orientation. No genomic duplications or unintended modifications were noted. Because the WGS confirmed we correctly generated the desired modification in this founder, we performed further validation of the model using this founder.

### Cre-mediated recombination of new Fzd2-flox allele results in Fzd2 knockout

To assess cre-mediated recombination of *loxP* sites (**Figure 4A**) in vitro, we cultured limb bud cells in micromass. This culture method was chosen due to the high expression of *Fzd2* mRNA in the developing limb according to ENCODE total RNA-seq data on E10.5 to E16.5 embryos [32], data not shown). Limb micromass cultures from wild-type or *Fzd2*^*tm1Vari*^ homozygous flox animals were infected with adenovirus containing Cre (Ad5CMV:Cre-eGFP) or control adenovirus (Ad5CMV:eGFP). Infection was confirmed by fluorescent imaging (**Figure 4B**). Genotyping by PCR amplification confirmed that a product was present at the anticipated size after cre-mediated recombination (**Figure 4C**). Sanger sequencing confirmed cre-mediated recombination resulted in the deletion of the Fzd2 coding sequence between the *loxP* sites, retaining sequence for one *loxP* site and XhoI and AgeI restriction sites (**Figure 4A**). Cell suspensions were stained using an antibody to FDZ2 and analyzed by flow cytometry. *Fzd2*^*tm1Vari*^ homozygous flox micromass cultures infected with Ad5CMV:Cre-eGFP showed a reduction in FZD2 expression compared to control, confirming cre-mediated deletion decreased FZD2 protein (**Figure 4D**).

To confirm *in vivo* cre-mediated recombination, we crossed female mice homozygous for CMV-cre (Tg/Tg) with male mice homozygous for the *Fzd2*^*tm1Vari*^ allele. CMV-cre was used to delete *Fzd2* in all tissues, including germ cells. Genotyping by PCR amplification confirmed that a product was present at the anticipated size for a gene knockout (KO) (**Figure 4E**). The wild-type (**Figure 4F**) or KO (**Figure 4G**) products were digested with XhoI or AgeI, resulting in expected product sizes. Based on the genotyping from 16 embryos generated from heterozygous (KO/+) crosses, we have not identified homozygous knockout animals (**Figure 4H**) as early as E12.5, consistent with the results from the IMPC (embryonic lethal prior to E9.5) [15]. Additional heterozygous crosses are needed to determine lethality at earlier embryonic stages.

## Discussion

Wnt signaling is critical for organ development and homeostasis. Many human diseases result from dysregulated Wnt signaling, making the pathway an important therapeutic target for diverse conditions, including osteoporosis, myocardial infarction, and cancer. Because there are 19 WNT ligands [33] that act through 10 Frizzled (FZD) receptors [34] and numerous associated co-receptors, therapeutic specificity is required in order to treat one disease without inadvertently causing debilitating side effects. These challenges are exacerbated by the fact that Wnts activate several signaling pathways, including the β-catenin (canonical) pathway [35, 36] and β-catenin-independent (non-canonical) pathways [37, 38]. Clinical trials to target Wnt signaling have confirmed these challenges. For example, Phase 1 clinical trials on cancer patients treated with Porcupine inhibitors, which impair all Wnt ligand secretion, or Vantictumab, which blocks signaling through 5 of the 10 FZDs, had to be halted due to severe osteoporosis as an unintended consequence [39]. Therefore, to create therapies that minimize harmful side effects, it is important to understand and target each Wnt pathway component’s unique physiologic function.

Studying one of the Frizzled receptors, Fzd2, poses a unique opportunity to better understand receptor specificity. Previous studies using knockout mouse models reported that Fzd2 had highly redundant physiological functions with Fzd1 and Fzd7 [13, 14]. Mice homozygous for this Fzd2 allele (*Fzd2*^*tm1*.*1Nat*^) were viable to birth, but half died neonatally due to cleft palate. Phenotypes were more dramatic when double mutants were generated. Specifically, double Fzd2 and Fzd7 mutant embryos died before gastrulation, and double Fzd1 and Fzd2 knockouts had fully penetrant cleft palate. However, based on our studies, we hypothesize that Fzd2 uniquely regulates early embryonic development without the functional redundancy with other Fzds previously reported. The studies which demonstrated functional redundancy between Fzd1, 2, and 7 were based on a knockout mouse model of Fzd2 (*Fzd2*^*tm1*.*1Nat*^) that retains the C-terminal region of coding sequence. While it might seem safe to assume that deletion of the N-terminal region which includes the first several transmembrane-spanning domains of a seven-transmembrane receptor would render it a null allele, we found the retained C-terminal sequence contained several potential in-frame start sites with strong Kozak sequences [30]. We created a plasmid directing expression of the remnant Fzd2 fragment starting from one of these methionines and found that it retained signaling capabilities when expressed in vitro. In further support of a unique role for Fzd2 in development, the IMPC generated a Fzd2 knockout mouse that, in contrast to the *Fzd2*^*tm1*.*1Nat*^ allele, deletes the entire coding sequence and is embryonic lethal prior to E9.5 [15]. Thus, it is likely that the true null phenotype is embryonic lethal early in development, making it impossible to determine specific functions of Fzd2 in any tissues formed after E9.5 in a global knockout animal.

We obtained a previously published Fzd2 flox model to determine the tissue-specific functions of Fzd2. The *Fzd2*^*tm1Eem*^ flox allele [16] was reported to develop lung cysts and loss of branching in embryonic lungs in *Shh-cre*;*Fzd2*^*tm1Eem*^ homozygotes. Given the association of FZD2 mutations with the rare human syndromes Autosomal Dominant Omodysplasia [8-11] and Robinow Syndrome (RS) [12], which present with craniofacial anomalies and limb reductions, we wanted to assess the phenotypes of mice lacking Fzd2 in these tissues. Interestingly, we saw dramatic phenotypes when we heterozygously deleted Fzd2 in limb precursor cells (*Prx1-cre* [40]) and cranial neural crest cells (*Wnt1-cre2* [41]) (data not shown). Interestingly, we generated global heterozygotes and homozygotes using *CMV-cre*, and these animals did not display any discernable phenotypes. These contradictory findings led us to believe there was a unique alteration in the *Fzd2*^*tm1Eem*^ allele. We sought to determine if this allele was a true null allele or if it altered FZD2 function in another way.

To determine if FZD2 protein expression was changed in the global mutants of the *Fzd2*^*tm1Eem*^ allele (*CMV-cre*; *Fzd2*^*tm1Eem*^), we used a new/non-commercial FZD2 antibody that specifically recognizes the N-terminal region of FZD2 and is validated for flow cytometry-based assays [27]. Using this antibody, we found that global mutants for the *Fzd2*^*tm1Eem*^ allele showed a modest reduction in FZD2, but not to the level of the *Fzd2*^*tm1*.*1Nat*^ homozygous mutants, which completely lack the Fzd2 N-terminus. This demonstrates an incomplete loss of FZD2 protein in the *Fzd2*^*tm1Eem*^ allele, suggesting the *Fzd2*^*tm1Eem*^ allele is not a correctly floxed model. Mice homozygous for the *Fzd2*^*tm1Eem*^ allele have significantly increased levels of *Fzd2* genomic DNA. WGS of the *Fzd2*^*tm1Eem*^ allele confirmed increased Fzd2 gene copy number.

To determine differences in the two copies of Fzd2, we used Sanger sequencing from PCR products amplifying sequence around the presumed 5’ or 3’ *loxP* sites. Sanger sequencing showed that one of the Fzd2 copies (Fzd2 #1) contains a WT 5’ sequence (lacks *loxP* site) and a 3’ sequence containing the *loxP/FRT* insertion. The other copy of Fzd2 (Fzd2 #2) contains a 5’ sequence with a *loxP* insertion and a 92bp deletion immediately upstream of the ATG start codon. Fzd2 #2 contains a WT 3’sequence with no *loxP* site. Based on these findings, we aimed to determine the result of Cre-mediated recombination on this allele.

We hypothesize that the two *loxP* sites are in opposite orientations leading to sequence inversion between these two *loxP* sites. We generated a “mutant” allele in vivo using *CMV-cre* and bred out the cre transgene to prevent further cre-mediated recombination. WGS of “mutant” mice did not show a loss of the region between the *loxP* sites, suggesting an inversion. If this was true, adding cre to “mutant” cells should result in a reversion to the “flox” allele. We collected limb buds from mutant embryos and cultured the cells in micromass. Exposure to adenovirus containing *CMV-cre* resulted in the restoration of the flox PCR product (data not shown). This reversion to the “flox” allele supports the hypothesis that cre-mediated recombination of the *Fzd2*^*tm1Eem*^ allele results in a continuous inversion. Thus, the *Fzd2*^*tm1Eem*^ allele is not a simple floxed allele, but, in fact, a complex genomic alteration that does not lead to a complete null for *Fzd2* upon exposure to Cre.

To better understand how Fzd2 regulates development, we generated a new Fzd2 flox model (*Fzd2*^*tm1Vari*^) by modifying recent approaches that enhance CRISPR-Cas9-mediated targeted editing. Based on the problems we noted with the previous mouse models of Fzd2 and because Fzd2 is a single exon gene and the whole gene is within a CpG island, we aimed to place *loxP* sites fully outside the Fzd2 locus but also avoid excessive elimination of genomic DNA that could unintentionally affect other gene expression after Cre-mediated deletion. We inserted the 5’ *loxP* in a region that is ∼2kb upstream of the ATG start codon lacking predicted transcription factor binding sites or promoter/enhancer marks. We placed the 3’ *loxP* site ∼1kb downstream of the 3’UTR to minimize effects on endogenous Fzd2 expression. We also chose to place the *loxP* sites in regions that would facilitate PCR-based genotyping since we noted difficulties in amplifying certain regions of the gene by PCR due to high or low GC content.

To facilitate the generation of a new Fzd2 flox model quickly and efficiently, we modified an approach recently published by the Rossant laboratory: two-cell (2C) homologous recombination (HR) CRISPR [7]. This method uses an mRNA that directs the expression of a Cas9-streptavidin fusion protein and further incorporates the use of biotinylated PCR templates to more accurately direct double-stranded cutting and facilitate the insertion of large modifications. Gu et al. also exploited the extended G2 phase of the cell cycle and the open chromatin environment at the two-cell stage to facilitate increased efficiency of genomic editing. Given our goal of the simultaneous insertion of two *loxP* sites, we made several modifications to this approach to potentially increase efficiency. To remove the step of converting Cas9 mRNA to protein in vivo after microinjection, we produced recombinant streptavidin tagged Cas (Cas9-mSA) protein for injection. Additionally, we formed ribonucleoprotein (RNP) complexes containing the Cas9-mSA, sgRNA, and biotinylated DNA repair template for each of the two *loxP* sites to increase efficiency and allow to target both *loxP* sites simultaneously.

Given our transgenic core’s extensive experience harvesting zygotes at the one-cell stage, we attempted two strategies for two-cell injections. The first strategy involved harvesting zygotes at the one-cell stage and culturing them until two-cell, at which point they were microinjected with RNP complexes and transferred to pseudopregnant females. The second strategy was to harvest zygotes at the two-cell stage and immediately microinject, followed by transfer. In our hands, we had higher success rates for the one-cell culturing method. Further work will establish which one of these methods is more advantageous.

Due to the nature of the two-cell injection, founders have the potential to be mosaic for the intended mutation. Backcrossing to C57BL/6J identifies founders that incorporated both *loxP* sites not only within the germline but also on the same allele. We further validated our model by extensive screening using Sanger sequencing, restriction digestions, and WGS. Given our experience with the unexpected and unintended issues noted with the *Fzd2*^*tm1Eem*^ allele, we highly encourage stringent validation of new mouse models using currently available methods. Because the cost of WGS has decreased dramatically in the last several years, it is now a cost-effective and arguably necessary tool to ensure scientific rigor.

We report here that we have successfully generated a floxed mouse strain where two *loxP* sites were simultaneously inserted around a region of interest with a single injection utilizing components of the CRISPR/Cas9 system. Our preliminary data suggest that global knockouts of Fzd2 (*CMV-Cre;Fzd2*^*tm1Vari*^) are early embryonic lethal, consistent with the IMPC knockout model. Future work will further define the timepoint in embryonic development at which the absence of Fzd2 leads to developmental arrest. We will also pursue studies using tissue-specific Cre drivers to study the role of Fzd2 in different developing structures. Altogether, we have successfully generated a new tool to study how Fzd2 functions *in vivo*, and the results of future studies will impact how we understand Wnt receptor specificity in development and disease.

## Acknowledgments

This work was funded by the Van Andel Institute. We would like to thank Edward Morrisey (University of Pennsylvania) and Ethan David Cohen (University of Rochester) for providing the *Fzd2*^*tm1Eem*^ mice and for discussions regarding the original design of the allele. We thank Jeremy Nathans (Johns Hopkins University) for providing us the *Fzd2*^*tm1*.*1Nat*^ mice. We would like to thank members of Janet Rossant’s laboratory (University of Toronto) for technical guidance regarding the two cell homologous recombination methods. Key members of the VAI Vivarium and Transgenics Core included Bryn Eagleson, Adam Rapp, Nicholas Getz, Brandon Bonnema, Audra Guikema, Tristan Kempston, Malista Powers, and Tina Schumaker. We thank the members of the Van Andel Institute Genomics Core and Flow Cytometry Core for assistance with whole-genome sequencing and flow cytometric analyses, respectively.

## Supplemental Figure Legends

**Supplemental Figure 1.**
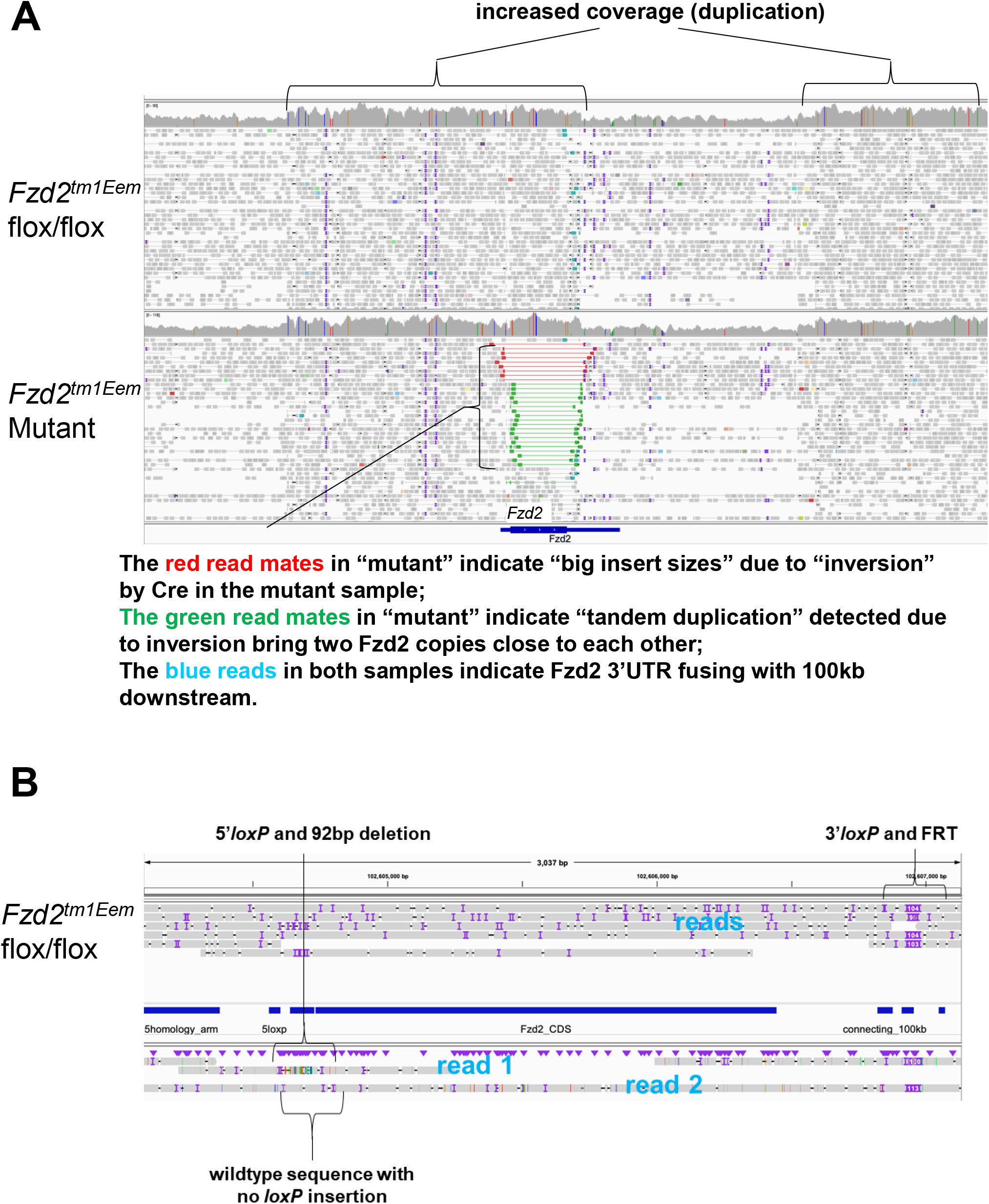
Whole-genome sequencing of *Fzd2*^*tm1Eem*^ alleles. A. Illumina sequencing of *Fzd2*^*tm1Eem*^ (homozygous flox “flox/flox” or homozygous global mutants “Mutant”). Reads were aligned to mm10 mouse reference genome for sequence and structural accuracy. The paired “big insert size” reads in red are possibly caused by Cre-driven inversion indicated in Figure 2A. Note no coverage reduction in the mutant sample. **B**. Nanopore (ONT) long-read sequencing was performed on gDNA from *Fzd2*^*tm1Eem*^ flox/flox animals. We identified two kinds of floxed Fzd2 alleles. One has a 5’*loxP* insertion along with a 92bp deletion downstream (read 1), while another *Fzd2*^*tm1Eem*^ allele does not have a 5’*loxP* or 92bp deletion but does have a 3’*loxP*/FRT insertion (read 2 and reads). We do not have a read that contains both 5’*loxP* insertion and 3’UTR region to confirm the nonexistence of a Fzd2 allele with both 5’ and 3’ *loxP* insertions, which is inferred by other evidence presented in the Discussion.

**Supplemental Figure 2.**
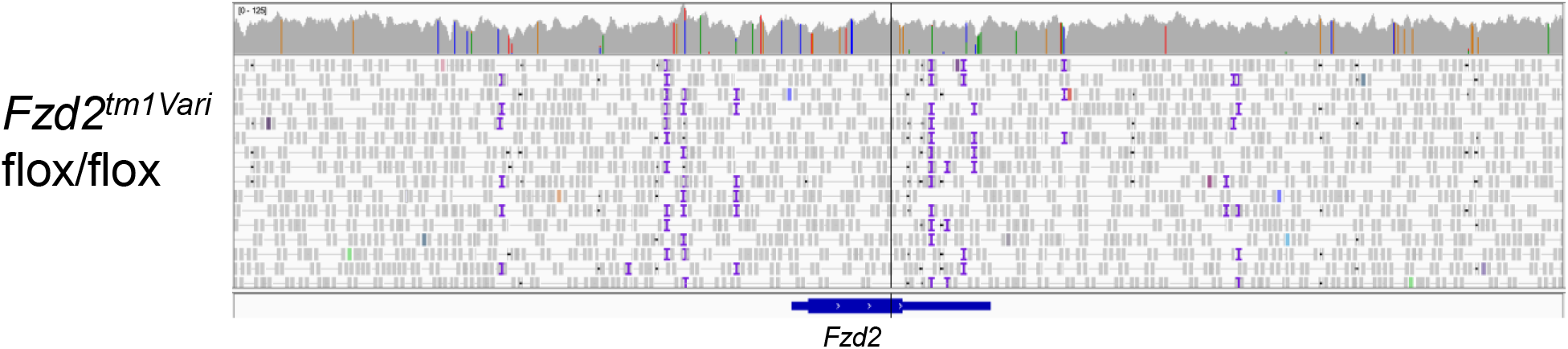
Whole-genome sequencing of *Fzd2*^*tmiVari*^ flox allele. Illumina sequencing of *Fzd2*^*tm1Vari*^ flox/flox allele. Reads were aligned to mm10 mouse reference genome for sequence and structural accuracy.

**Supplemental Figure 3.**
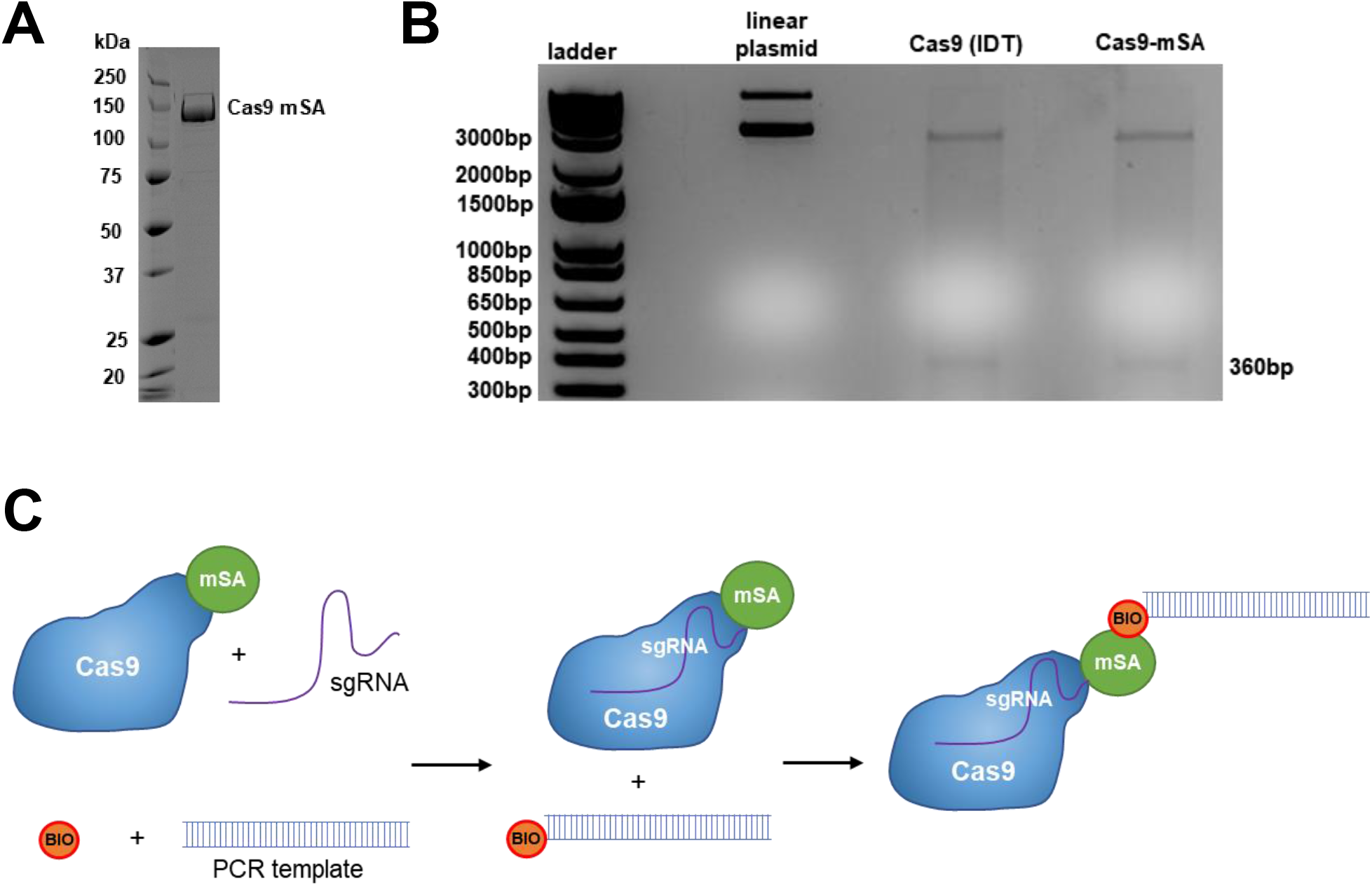
Cas9-mSA recombinant protein. A. SDS page after size exclusion column. **B**. DNA cleavage activity of Cas9-mSA endonuclease was assayed on linearized plasmid DNA containing eGFP (LentiCMVtight-GFP) and produced a cut product of the expected size (360bp). **C**. Ribonucleoprotein (RNP) complexes containing Cas9-mSA fusion protein complexed with specific sgRNA (sgRNA1 or sgRNA 2 details in the Methods section and Figure 3A) were covalently linked to biotinylated PCR templates.

**Supplemental Figure 4.**
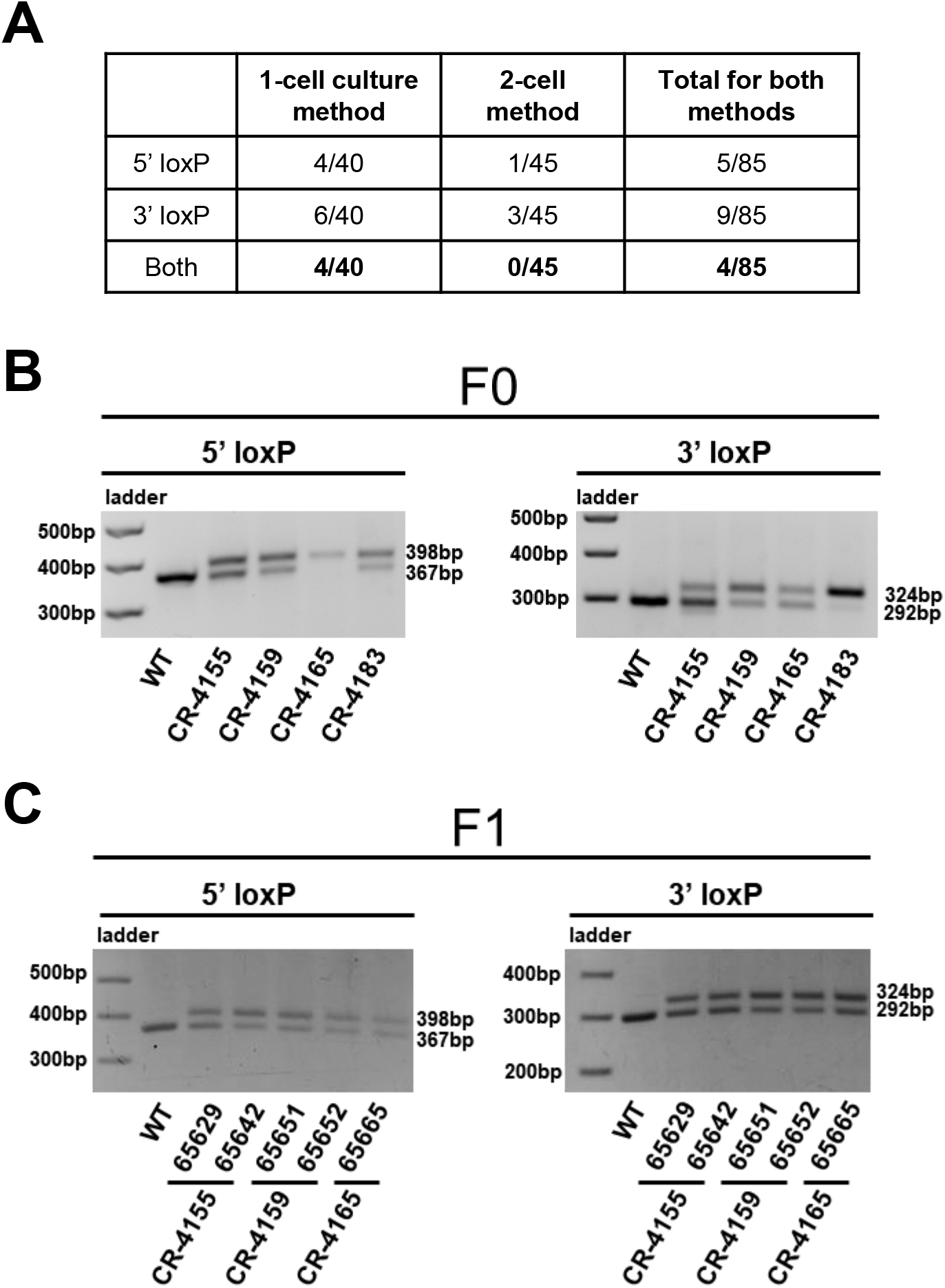
Founder information. A. Table of resulting founders from the 1-cell culture or 2-cell methods. **B**. PCR-based genotyping results of F0 offspring. **C**. PCR-based genotyping results of F1 progeny.

## References

1. Orban, P.C., D. Chui, and J.D. Marth, Tissue- and site-specific DNA recombination in transgenic mice. Proc Natl Acad Sci U S A, 1992. 89(15): p. 6861–5.

2. Schwenk, F., U. Baron, and K. Rajewsky, A cre-transgenic mouse strain for the ubiquitous deletion of loxP-flanked gene segments including deletion in germ cells. Nucleic Acids Res, 1995. 23(24): p. 5080–1.

3. Yang, H., et al., One-step generation of mice carrying reporter and conditional alleles by CRISPR/Cas-mediated genome engineering. Cell, 2013. 154(6): p. 1370–9.

4. Quadros, R.M., et al., Easi-CRISPR: a robust method for one-step generation of mice carrying conditional and insertion alleles using long ssDNA donors and CRISPR ribonucleoproteins. Genome Biology, 2017. 18(1): p. 92.

5. Liu, Y., et al., Generation of Conditional Knockout Mice by Sequential Insertion of Two loxP Sites In Cis Using CRISPR/Cas9 and Single-Stranded DNA Oligonucleotides. Methods Mol Biol, 2019. 1874: p. 191–210.

6. Horii, T., et al., Efficient generation of conditional knockout mice via sequential introduction of lox sites. Sci Rep, 2017. 7(1): p. 7891.

7. Gu, B., E. Posfai, and J. Rossant, Efficient generation of targeted large insertions by microinjection into two-cell-stage mouse embryos. Nat Biotechnol, 2018. 36(7): p. 632–637.

8. Nagasaki, K., et al., Nonsense mutations in FZD2 cause autosomal-dominant omodysplasia: Robinow syndrome-like phenotypes. Am J Med Genet A, 2018. 176(3): p. 739–742.

9. Saal, H.M., et al., A mutation in FRIZZLED2 impairs Wnt signaling and causes autosomal dominant omodysplasia. Hum Mol Genet, 2015. 24(12): p. 3399–409.

10. Turkmen, S., et al., A Novel de novo FZD2 Mutation in a Patient with Autosomal Dominant Omodysplasia. Mol Syndromol, 2017. 8(6): p. 318–324.

11. Warren, H.E., et al., Two unrelated patients with autosomal dominant omodysplasia and FRIZZLED2 mutations. Clin Case Rep, 2018. 6(11): p. 2252–2255.

12. White, J.J., et al., WNT Signaling Perturbations Underlie the Genetic Heterogeneity of Robinow Syndrome. Am J Hum Genet, 2018. 102(1): p. 27–43.

13. Yu, H., et al., Frizzled 1 and frizzled 2 genes function in palate, ventricular septum and neural tube closure: general implications for tissue fusion processes. Development, 2010. 137(21): p. 3707–17.

14. Yu, H., et al., Frizzled 2 and frizzled 7 function redundantly in convergent extension and closure of the ventricular septum and palate: evidence for a network of interacting genes. Development, 2012. 139(23): p. 4383–94.

15. Dickinson, M.E., et al., High-throughput discovery of novel developmental phenotypes. Nature, 2016. 537(7621): p. 508–514.

16. Kadzik, R.S., et al., Wnt ligand/Frizzled 2 receptor signaling regulates tube shape and branch-point formation in the lung through control of epithelial cell shape. Proc Natl Acad Sci U S A, 2014. 111(34): p. 12444–9.

17. Moreno-Mateos, M.A., et al., CRISPRscan: designing highly efficient sgRNAs for CRISPR-Cas9 targeting in vivo. Nat Methods, 2015. 12(10): p. 982–8.

18. Savic, N., et al., Covalent linkage of the DNA repair template to the CRISPR-Cas9 nuclease enhances homology-directed repair. Elife, 2018. 7.

19. Savić, N., et al., In vitro Generation of CRISPR-Cas9 Complexes with Covalently Bound Repair Templates for Genome Editing in Mammalian Cells. Bio Protoc, 2019. 9(1).

20. Mali, P., et al., RNA-guided human genome engineering via Cas9. Science, 2013. 339(6121): p. 823–6.

21. Mehravar, M., et al., In Vitro Pre-validation of Gene Editing by CRISPR/Cas9 Ribonucleoprotein. Avicenna J Med Biotechnol, 2019. 11(3): p. 259–263.

22. Harms, D.W., et al., Mouse Genome Editing Using the CRISPR/Cas System. Curr Protoc Hum Genet, 2014. 83: p. 15.7.1-27.

23. Iezaki, T., et al., Cartilage Induction from Mouse Mesenchymal Stem Cells in Highdensity Micromass Culture. Bio-protocol, 2019. 9(1): p. e3133.

24. Underhill, T.M., H.J. Dranse, and L.M. Hoffman, Analysis of Chondrogenesis Using Micromass Cultures of Limb Mesenchyme, in Skeletal Development and Repair: Methods and Protocols, M.J. Hilton, Editor. 2014, Humana Press: Totowa, NJ. p. 251–265.

25. Nusspaumer, G., et al., Ontogenic Identification and Analysis of Mesenchymal Stromal Cell Populations during Mouse Limb and Long Bone Development. Stem Cell Reports, 2017. 9(4): p. 1124–1138.

26. Reinhardt, R., et al., Molecular signatures identify immature mesenchymal progenitors in early mouse limb buds that respond differentially to morphogen signaling. Development, 2019. 146(10): p. dev173328.

27. Tao, Y., et al., Tailored tetravalent antibodies potently and specifically activate Wnt/Frizzled pathways in cells, organoids and mice. Elife, 2019. 8.

28. D’Haene, B., J. Vandesompele, and J. Hellemans, Accurate and objective copy number profiling using real-time quantitative PCR. Methods, 2010. 50(4): p. 262–70.

29. Eubelen, M., et al., A molecular mechanism for Wnt ligand-specific signaling. Science, 2018. 361(6403).

30. Kozak, M., An analysis of 5’-noncoding sequences from 699 vertebrate messenger RNAs. Nucleic Acids Res, 1987. 15(20): p. 8125–48.

31. de Lau, W., et al., The R-spondin/Lgr5/Rnf43 module: regulator of Wnt signal strength. Genes Dev, 2014. 28(4): p. 305–16.

32. Davis, C.A., et al., The Encyclopedia of DNA elements (ENCODE): data portal update. Nucleic Acids Res, 2018. 46(D1): p. xD794–d801.

33. Mikels, A.J. and R. Nusse, Wnts as ligands: processing, secretion and reception. Oncogene, 2006. 25(57): p. 7461–8.

34. Wang, Y., et al., Frizzled Receptors in Development and Disease. Curr Top Dev Biol, 2016. 117: p. 113–39.

35. Nusse, R. and H. Clevers, Wnt/β-Catenin Signaling, Disease, and Emerging Therapeutic Modalities. Cell, 2017. 169(6): p. 985–999.

36. Daulat, A.M. and J.P. Borg, Wnt/Planar Cell Polarity Signaling: New Opportunities for Cancer Treatment. Trends Cancer, 2017. 3(2): p. 113–125.

37. Veeman, M.T., J.D. Axelrod, and R.T. Moon, A second canon. Functions and mechanisms of beta-catenin-independent Wnt signaling. Dev Cell, 2003. 5(3): p. 367–77.

38. Romereim, S.M. and A.T. Dudley, Cell polarity: The missing link in skeletal morphogenesis? Organogenesis, 2011. 7(3): p. 217–28.

39. Mirabelli, C.K., et al., Perspectives on the role of Wnt biology in cancer. Sci Signal, 2019. 12(589).

40. Logan, M., et al., Expression of Cre Recombinase in the developing mouse limb bud driven by a Prxl enhancer. Genesis, 2002. 33(2): p. 77–80.

41. Lewis, A.E., et al., The widely used Wnt1-Cre transgene causes developmental phenotypes by ectopic activation of Wnt signaling. Dev Biol, 2013. 379(2): p. 229–34.

